# Unsupervised Deep Learning for Cryo-EM/ET Particle Classification

**DOI:** 10.64898/2026.11.12.766036

**Authors:** Mostofa Rafid Uddin, Seonghui Min, Qifeng Wu, Xueying Zhan, Muyuan Chen, Min Xu

**Author notes:** Equal contributions (Mostofa Rafid Uddin, Seonghui Min). {, }.

## Abstract

Cryogenic electron microscopy (cryo-EM) is widely used to determine protein structures from purified samples. With cryogenic electron tomography (cryo-ET), it is possible to image thin sections of cells and directly visualize diverse proteins in their native environment. However, separating the massive mixture of different types of proteins inside cells remains challenging, especially without human intervention. In this work, we present an unsupervised deep learning framework for classifying cryo-EM/ET particle images that jointly optimizes for pose and structure through an autoencoder-based multi-stage pipeline. On single-particle cryo-EM datasets, our framework can capture the continuous variation in particle views and separate images of real particles from contamination. Using *in silico* mixtures of multiple types of proteins, we demonstrated our method’s capability of separating the protein classes without supervision. Finally in a real cryo-ET dataset of algal chloroplast, our deep learning model successfully detected the four main protein complexes of photosynthesis on native thylakoid membranes. Taken together, these results demonstrate that our framework offers a scalable, annotation-free solution for robust structural discovery across diverse cryo-EM and cryo-ET applications.

## Introduction

Single-particle cryogenic-electron microscopy (cryo-EM) has become the dominant technique for determining macromolecular structures at near-atomic resolution [1, 2]. While structural biology research typically requires isolating and purifying homogeneous macromolecules [3], with advances in cryogenic electron tomography (cryo-ET) [4] and sample preparation techniques [5], it has become increasingly routine to image thin sections of individual cells and study protein structures in their native environment.

However, current in situ cryo-ET studies are primarily limited to a few well-characterized proteins, such as ribosomes and microtubules, which are large and abundant in cells. A key challenge in studying more diverse proteins is differentiating them within crowded cellular spaces amid noisy data. With a mixture of many different proteins, it becomes nearly impossible even for human experts to separate different protein densities and select particles of the same type of protein for high-resolution averaging.

In some cases, template matching can be used to identify certain types of proteins from cellular tomograms [6, 7]. However, template-based methods require existing structures of the target protein and the assumption that the in situ structures sufficiently resemble the in vitro ones. Additionally, iterative 2D and/or 3D classification has been used to separate the different types of proteins from mixtures [8, 9], but these methods require input from human experts to select the “good” classes, which can potentially introduce unwanted bias.

More recently, learning-based approaches have been developed to analyze structural heterogeneity in cryo-EM and cryo-ET data [10, 11, 12, 13]. Other unsupervised methods have focused on identifying structurally distinct macromolecules directly from cellular subtomograms [14, 15]. In this work, we developed an unsupervised deep-learning-based approach for classifying cryo-EM/ET particle images in a fully automated, data-driven manner.

At a high level, our method uses an autoencoder network trained with pose augmentation and a “winner-take-all” reconstruction loss to disentangle protein structure from its in-plane orientation. To learn structural representations in a coarse-to-fine manner, a three-stage training strategy was adopted to optimize the autoencoder network. After training, the particles are embedded in a low-dimensional latent space that depicts both the viewing angle and the 3D structural features of the particles, allowing the separation of the different types of particles.

We validated our pipeline through experiments on multiple cryo-EM/ET datasets, demonstrating its ability to capture continuous variation in particle views and perform automated particle curation in single-particle cryo-EM data, distinguish particle classes in *in silico* mixtures, and separate different types of macromolecular complexes in cellular cryo-ET data.

## Results

### Overview

Our method employs an autoencoder framework to learn latent representations from cryo-EM particle images while estimating their in-plane poses. For each particle image, the encoder maps an input image to a latent representation, and the decoder generates a denoised 2D structural template. As in a variational autoencoder [16], during training, a small random variable is added to the encoder output to ensure the smoothness of the latent space. To address the random in-plane rotation of the particles, a “winner-take-all” reconstruction loss [17, 18, 19, 20] was used in the training process. Specifically, multiple copies of each input image, rotated by a series of angles, are input to the encoder, and the initial loss function is calculated by comparing the decoder outputs to the corresponding rotated inputs. The “winner-take-all” loss is then computed by taking the minimum of the loss values from all rotated input images. Therefore, the autoencoder only needs to learn one correct representation of all in-plane rotated versions of the same particle image, without having to encode the exact rotation angle of the input image internally [17, 21, 15]. The “winner-take-all” loss reduces the burden of the deep learning model and greatly speeds up training convergence in our experiments. Additionally, while not directly encoded, the in-plane pose of each particle can still be computed, by simply taking the arguments of the minima of the initial loss values from the rotated inputs. The per-particle poses are stored outside the autoencoder model, and updated every epoch.

To mimic a typical alignment procedure of cryo-EM images [22], the training process is divided into three sequential stages (Fig. 1), each dealing with finer angular steps of rotation angles and higher-resolution information in the images. The same autoencoder model is used throughout the stages, with each later stage initialized using the weights learned in the preceding stage. Stage 1 uses down-sampled particle images and a coarsely sampled exhaustive alignment grid. Starting from stage 2, we take into account the in-plane poses determined previously, and only rotate the input images to a smaller range of angles with finer sampling around the existing poses. Higher-resolution images are also used in the later training stages to improve the alignment accuracy, and an additional boosting layer is introduced at stage 3 to better capture the high-resolution information.

**Figure 1.**
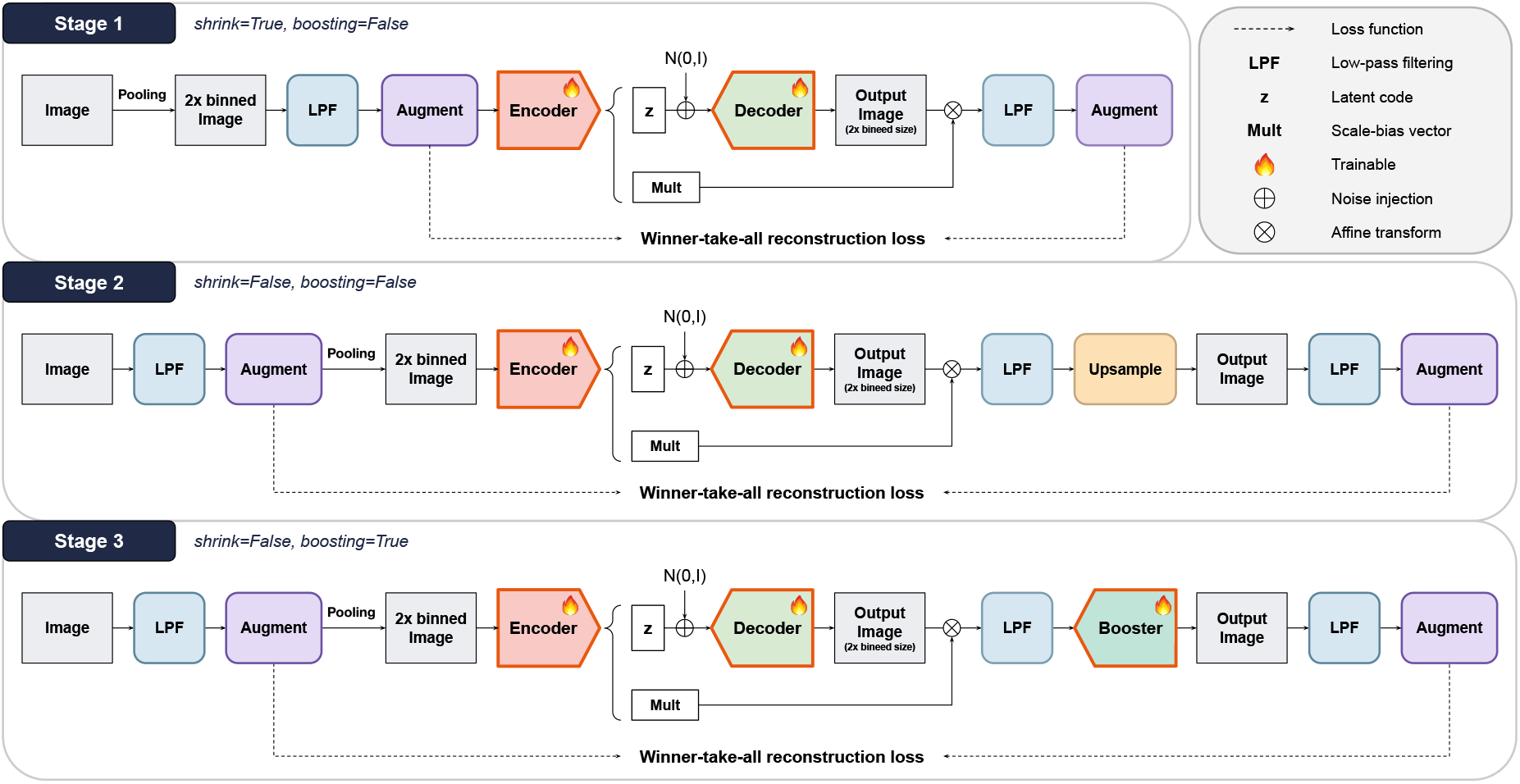
Overview of our unsupervised training pipeline. Stage 1 uses particle images and reconstruction targets down-sampled by a factor of 2; Stages 2 and 3 operate at full resolution. In Stage 2, the decoder output is upsampled to full resolution, while Stage 3 introduces a trainable booster decoder to generate the full-resolution output. Across all stages, the encoder produces a latent code *z* and a scale-bias vector, Gaussian noise is injected into *z*, and the resulting reconstruction output is compared with the corresponding target using the winner-take-all reconstruction loss.

After training, the learned representations, reconstruction losses, and decoded templates are used according to the analysis objective to represent particle-view variation, group particles, and guide the selection of particle subsets for downstream analysis.

### Experiments on single-particle cryo-EM datasets

We first evaluated our method on single-particle cryo-EM datasets. Starting from a simple example, we used a subset of 40,000 TRPV1 particles from EMPIAR-10059. The subset of particles was curated with existing methods [22] to exclude most non-particle images. After training, the model successfully organized the particles according to their projection angle. Neighboring regions in the latent space represent similar views and gradual transitions from side to top views of the protein can be seen across the latent space (Fig. 2A). The trained model correctly captured the protein structures and their poses in input images. Upon visual inspection, the decoder output matches the input image well, and subtracting the decoder output from the input leaves only the background noise. Interestingly, not only particles of the same projection angle are aligned by the autoencoder, particles of different views also appear visually aligned, likely due to the smoothness constraint of the latent space.

**Figure 2.**
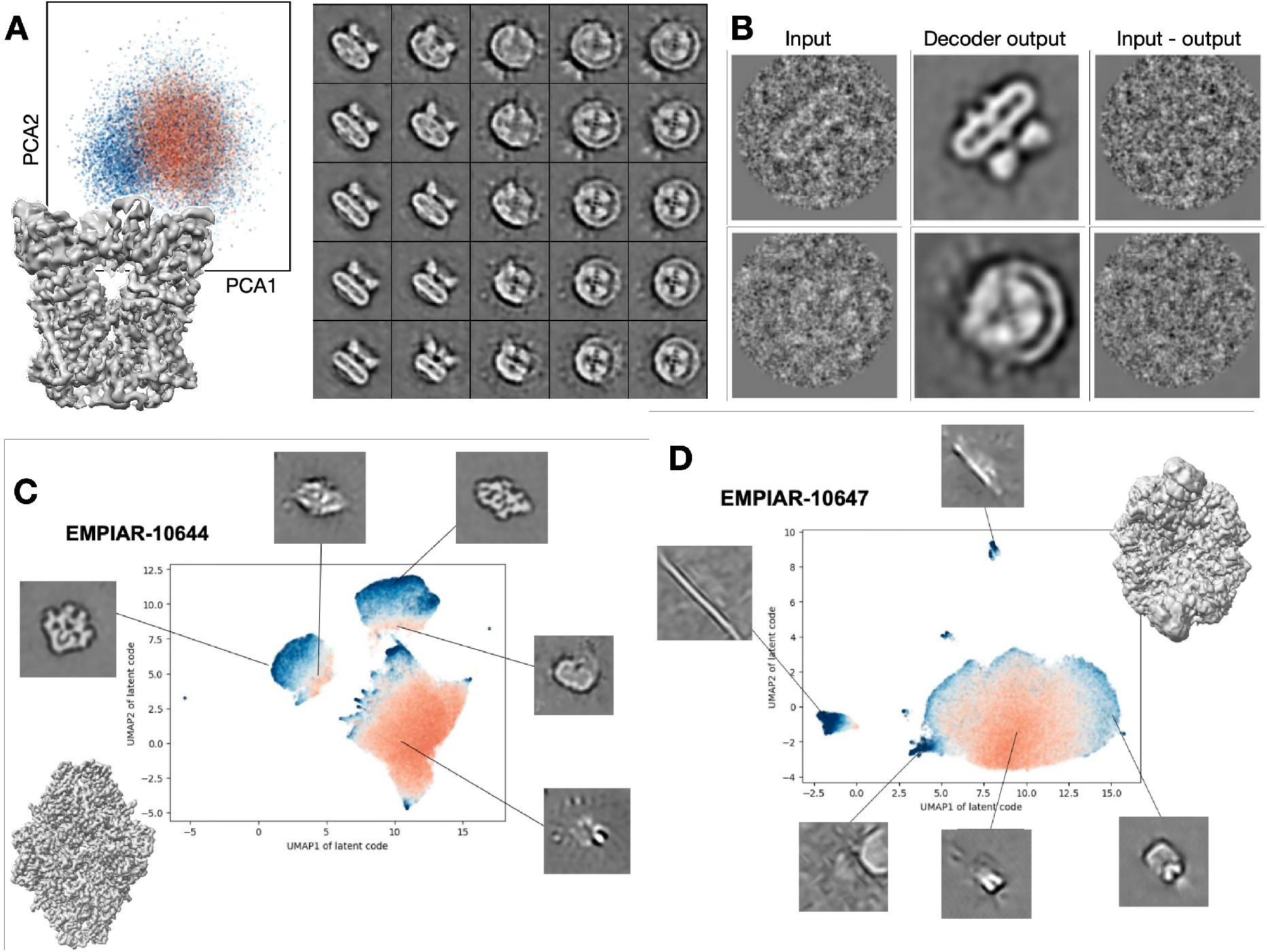
Latent-space representation of particle-view variation and automated particle curation in single-particle cryo-EM datasets. For TRPV1 **(A)**, the learned latent-space representation and representative decoded images sampled across the latent space illustrate continuous variation in particle views. **(B)** Examples of input images, the corresponding decoded images, and their residuals (input minus decoded image) show that particle structures are captured in the reconstructions, whereas the residuals contain primarily background noise. For *β*-galactosidase **(C)** and PKM2 **(D)**, the two-dimensional embeddings and representative decoded images show the separation of high-quality particles from low-quality or background images. In **(A), (C), and (D)**, points are colored by reconstruction loss on a continuous scale, ranging from blue for lower loss to red for higher loss.

**Figure 3.**
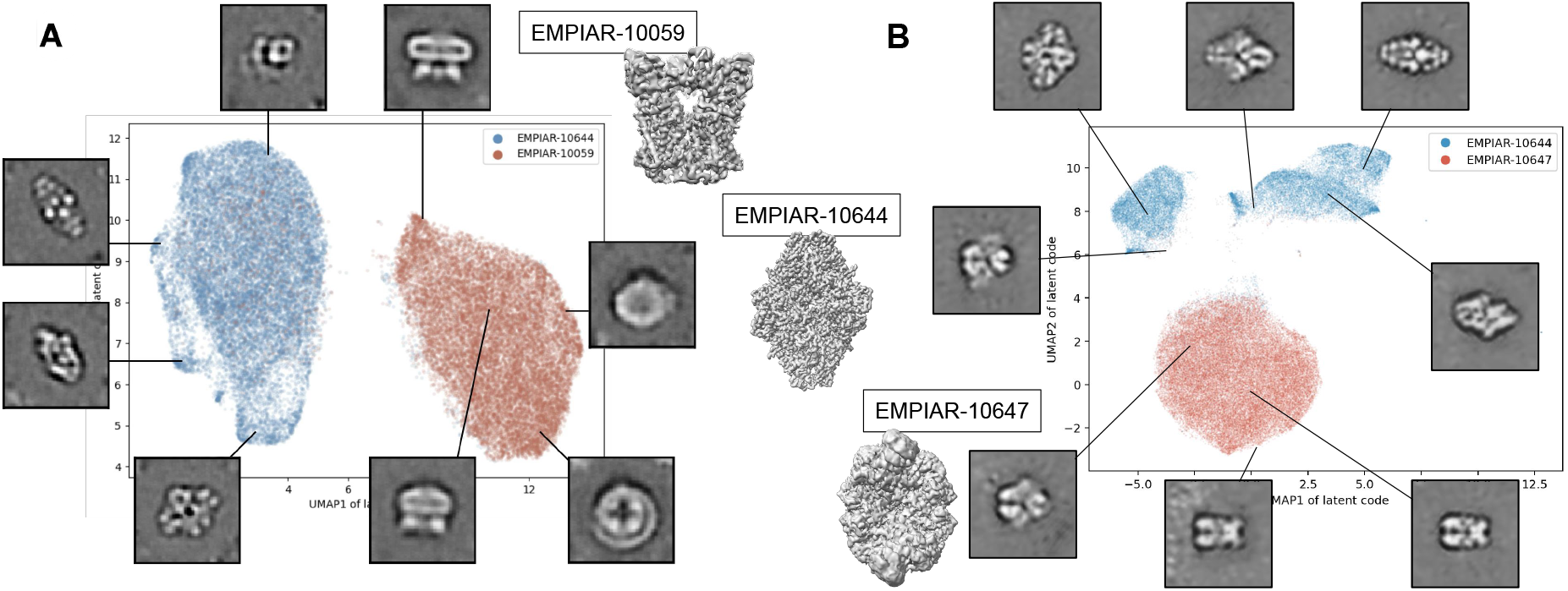
Classification of distinct particle classes in *in silico* mixtures of single-particle cryo-EM images. Mixtures of TRPV1 and *β*-galactosidase **(A)** and of *β*-galactosidase and PKM2 **(B)** are visualized in the learned latent space together with representative decoded images. The *β*-galactosidase– PKM2 mixture represents a more challenging case since the two complexes are more structurally similar. In both panels, points are colored by particle class, with the two classes shown in blue and red.

Next, we tested the method on full cryo-EM datasets to evaluate its ability to separate protein images from contamination and perform automated particle curation. Using 97,163 particle images automatically selected from a public *β*-galactosidase dataset (EMPIAR-10644), we showed that the autoencoder can separate images of proteins from those of non-protein features such as carbon edges and ice contamination. It is also worth noting that due to preferred orientations, images of *β*-galactosidase form two clusters as there are no transition views that connect them. Through simple clustering of the latent space, the method automatically identified 33,097 “good” particle images, which were subsequently used for cryoSPARC abinitio reconstruction. This yielded a 3D structure of *β*-galactosidase at 2.55Å resolution, compared with 2.60Å resolution when all particle images were used, suggesting that the method can successfully remove “junk” particles and produce a more homogeneous particle population.

Similarly, on a public dataset of the PKM2 complex from EMPIAR-10647, the autoencoder mapped images of actual particles and contamination to different clusters in the latent space. Based on their latent-space distribution, the method selected 51,128 of 84,500 particles, which yielded a 3D reconstruction at 2.91Å resolution, compared with 2.98Å when all 84,500 particles were used.

### Experiments on *in silico* cryo-EM particle mixtures

After evaluating our method on individual single-particle cryo-EM datasets, we next tested whether it could distinguish different structures in *in silico* mixtures of single-particle cryo-EM images.

To this end, we first created a mixture of *β*-galactosidase particle images from EMPIAR-10644 and

TRPV1 complex particle images from EMPIAR-10059, two proteins with substantial structural differences. The two datasets were resampled to the same pixel size and were both low-pass filtered to 5Å. When applied to this mixture dataset, our pipeline successfully mapped the particles of the two proteins to two distinct clusters in the latent space. Simple clustering in the latent space separated *β*-galactosidase from TRPV1 complexes, achieving a classification accuracy of 98.6%.

We then created another mixture dataset of *β*-galactosidase from EMPIAR-10644 and PKM2 complex from EMPIAR-10647, two proteins with greater structural similarity (both are soluble and have D2 symmetry). With the same protocol, the autoencoder produced a latent space that clearly separated the two datasets, achieving a classification accuracy of 96.7%. Notably, due to the preferred orientation, particles of *β*-galactosidase were separated into two disconnected clusters. However, the distance between the two clusters of the same protein was still smaller than the distance between the two types of proteins, making the classification possible.

After classifying particles by their positions in the latent space, a 3D structure of each class can be obtained through ab initio reconstruction from the corresponding 2D particles. The reconstructed 3D structures resemble those generated from the individual cryo-EM datasets prior to the *in silico* mixing.

### Experiments on cellular cryo-ET subtomograms

Upon successfully distinguishing particle classes in *in silico* mixtures of single-particle cryo-EM images, we demonstrated our method’s capability of distinguishing distinct macromolecular complexes in real *in situ* cellular cryo-ET data.

To this end, we used a cryo-ET dataset of *Chlamydomonas reinhardtii* publicly available in EMPIAR-11830, and focused on the protein complexes embedded on thylakoid membranes inside the chloroplast.

Compared to the mixture of potentially hundreds of thousands of different soluble proteins inside the cytosol, the thylakoid membrane provides a relatively simple example. It is already known that there are mainly four membrane proteins in the thylakoids [23], which facilitate photosynthesis in the chloroplast, and their structures have been characterized in vitro [24, 25, 26, 27].

To identify the photosynthesis components, we first selected particles containing protein-like densities inside the chloroplast. This was done semi-automatically using a previously established protocol [28, 29], and the neural network for particle picking was trained with sample particle images that had some additional density protruding from a piece of membrane. To simplify the problem, particles on the inner and outer chloroplast membranes were manually excluded.

After particle selection, CTF-corrected subtomograms were extracted from the tilt series, and cryo-EM-like images were generated by projecting the subvolumes along the depth (*z*) axis. The 2D images were then input to the autoencoder, which mapped them to a 2D latent space. The decoder outputs were well aligned, and clear densities of proteins were visible on top of the membranes (Fig. 4B). In the latent space, the particles were roughly divided into two groups, one with a single membrane, and the other with stacked membranes. Unlike the *in silico* mixture datasets, here the particles did not form clearly separable clusters.

**Figure 4.**
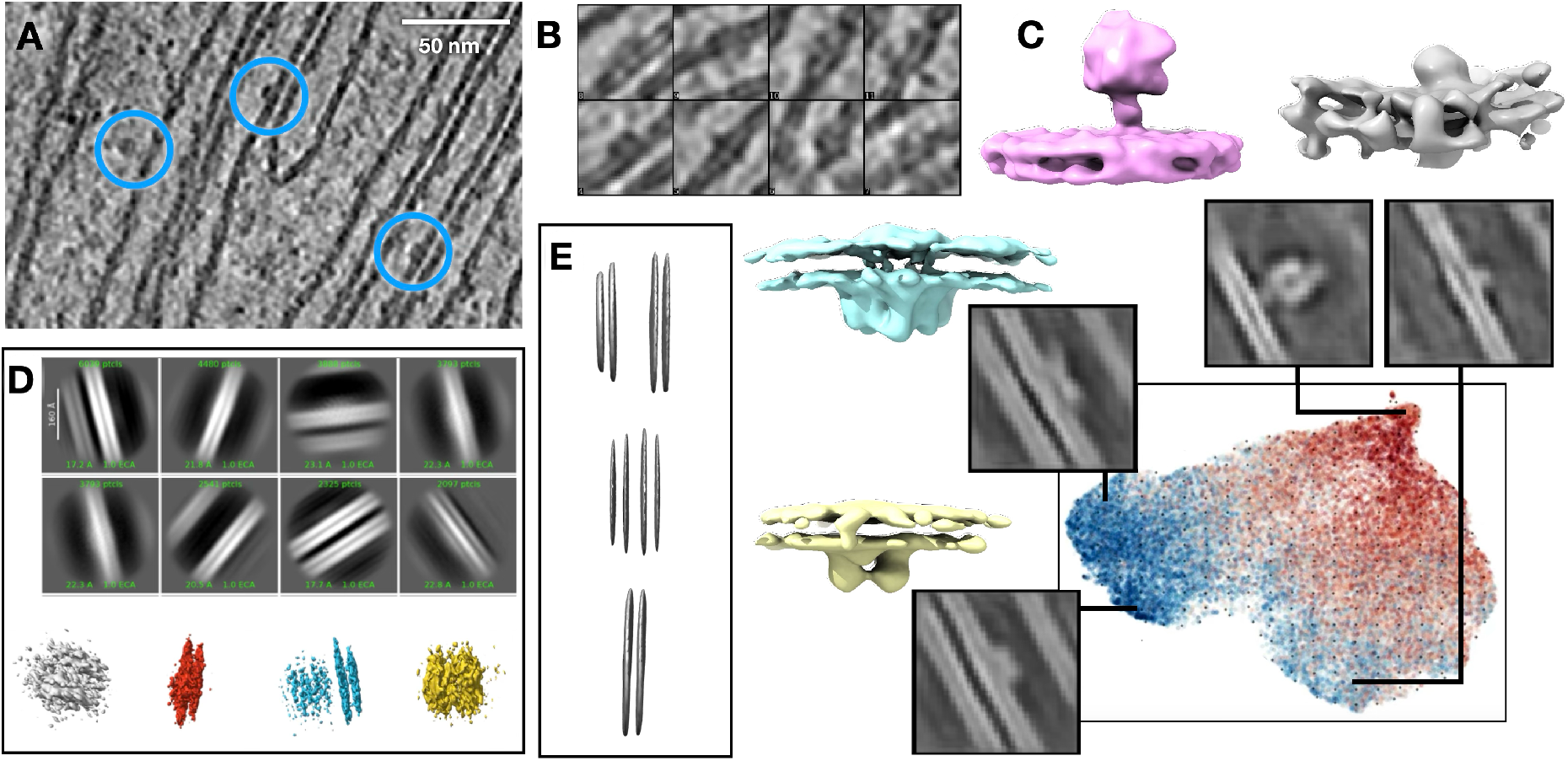
Identification of distinct macromolecular complexes in cellular cryo-ET data from *Chlamydomonas reinhardtii*. **(A)** Representative thylakoid membrane region in the tomogram, with example particle locations highlighted. **(B)** Example decoder outputs showing membrane-associated particle structures. **(C)** Using our method, the latent-space representation and representative decoded images reveal distinct membrane-associated structural patterns, with representative 3D reconstructions obtained from the corresponding particle subsets. Points in the latent-space representation are colored by reconstruction loss on a continuous scale, ranging from blue for lower loss to red for higher loss. **(D)** cryoSPARC 2D classification produced class averages dominated by membrane features; representative 3D volumes reconstructed from the corresponding particle subsets are shown below. **(E)** Representative 3D classes obtained using RELION 3D classification.

Likely due to the complexity of the *in situ* dataset, the space between clusters was occupied by what can be described as “bad” particles, including bare membranes with little or no protein density protrusion, and highly tilted particles that did not have sufficient membrane features for alignment.

Based on the features of the decoder output, we segmented the “good” particles of the latent space into four clusters, and generated an *ab initio* 3D reconstruction for each class using the corresponding 3D subtomograms [29]. In the 3D structures, clear protein densities could be identified on the thylakoid membranes, and their shapes generally resembled the four known thylakoid membrane proteins participating in photosynthesis.

For comparison, we also applied the classical 2D classification [30] to the projected 2D thylakoid membrane protein particles, as well as 3D classification [31] to the corresponding 3D subtomograms. Both approaches were able to resolve the membranes in the averaged structures and distinguish the particles with single or stacked membranes. However, despite our best efforts, we were unable to recover the densities of protein complexes expected to be present on the thylakoid membranes using the traditional approaches.

## Discussion

In this paper, we presented an unsupervised approach that maps cryo-EM/ET particle images to a low-dimensional latent space based on their structures, allowing the separation of different types of particles. Using an autoencoder with pose augmentation and a “winner-take-all” loss, our method can capture the continuous transitions in particle projection angles, so that particles that share the same 3D structure are attracted to the same latent space cluster. This enabled automated particle curation, the separation of particle classes from in silico mixtures, and the identification of distinct macromolecular complexes from in situ cryo-ET datasets.

Unlike most heterogeneity analysis techniques [10, 11, 32], the model requires no prior information about the pose or structure of the particles and can separate particles of completely different proteins that do not have a common core for alignment. Additionally, our method does not involve pre-training on existing datasets [33], avoiding the potential issue of poor cross-domain inference performance, particularly for in situ cryo-ET data.

Building on our previous unsupervised cryo-ET protein identification models [14, 15], the current method introduced a number of simplifications that have surprisingly improved its performance. Compared with our previous work and other existing methods [21], one key simplification is the removal of in-plane pose encoding within the encoder. By replacing the pose encoding with the “winner-take-all” loss, the training process becomes much smoother, and the model can robustly converge even for complex and noisy datasets.

Compared to methods that include a separate pose search parallel to conformation encoding [34], the “winner-take-all” loss ensures that pose and shape estimation are still integrated within a differentiable network, making it possible to perform shape-based particle separation and pose alignment simultaneously within the model training process.

The next simplification here is to separate the shape-based particle clustering step from the 3D reconstruction. While most deep-learning-based methods for cryo-EM/ET particle classification learn to encode some type of 3D representation of protein structures within the model [10, 11, 17], to our surprise, it turns out that 2D particles of the same protein still form clusters in latent space by themselves, without an explicit representation of 3D structures within the model. Admittedly, this requires a smooth transition between the different projection angles of the same 3D structure among the particles, which can become a challenge for single-particle cryo-EM datasets of proteins that have disconnected clusters of preferred views. However, since the main scenario involving particle mixtures occurs within the cellular environment, which is generally free from air-water interface effects [35], the requirement for smooth view transitions should be easier to meet. Additionally, for both cryo-EM and cryo-ET datasets, as long as particles of different types are reasonably separated, existing ab initio reconstruction algorithms [30, 29] can easily generate 3D structures from a cluster of particles, so the encoding of 3D structures within the deep learning model becomes even less necessary.

Finally, we opt to only use z-axis projection images of the subtomograms for the analysis of cryo-ET datasets. This approach was initially designed to maintain consistency between the analysis of cryo-EM and cryo-ET datasets and was implemented to be expandable so that multiple sub-tilt images could potentially be used as input to capture 3D information. However, in our experiments, the introduction of additional per-particle projection angles did not improve model performance. This might be because, for two subtomograms with the same z-projection image at two different in-plane rotation angles, their tilted views can differ drastically because their tilt axes would differ. The in-plane pose augmentation and the “winner-take-all” loss do not take this into account, which can end up separating the subtomograms into different clusters.

While these simplifications improve model training convergence and performance in our experiments, they also bring a number of limitations. In addition to the challenge of disconnected preferred orientations mentioned above, the separation of clusters in the latent space can also be corrupted by the non-particle images connecting view angle clusters. As shown in the examples of photosystems, “bad” particle images can sometimes serve as smooth transitions between clusters of different types of proteins, making the latent space less separable. While it is still possible to select particles based on their locations in the latent space and the corresponding decoder output, the process is more challenging, especially when there is insufficient prior knowledge about the system. Potentially, this issue can be solved by introducing an extra round of preprocessing to remove “bad” particles from the dataset. However, the additional step may also introduce more human bias as the decision of which particles are “bad” is often subjective.

Further development is still needed to achieve the ultimate goal of visual proteomics, i.e., distinguishing many different types of proteins in a crowded in situ environment. First, the current method only works on extracted particles and assumes there is only one protein within the particle image. In a system of proteins with widely varying sizes, the assumption no longer holds true. For example, an image large enough to contain one ribosome can contain multiple smaller enzymes, making it impossible to map the particle image to a single point in the latent space. One solution is to take an iterative approach [8] that starts by distinguishing the larger proteins and then excludes them from the particle selection process to search for smaller ones. Second, our model has currently been tested only on mixtures of 2-4 different proteins. A more complex neural network architecture may be required to capture more complex systems, and more data and computational resources may be needed for the development of such networks. Finally, although the current model performs well on our example datasets without explicit input of 3D information, it would still be advantageous to design an architecture that introduces 3D information to the model without sacrificing training convergence.

## 1 Method

### 1.1 Problem Setting

In cellular cryo-ET or cryo-EM datasets with multiple structural classes, the presence of a set of distinct 3D structure templates per structural class is assumed. Each 2D image in the cryo-EM dataset is generated from any one of these 3D template structures using a series of geometric and imaging operations. A mathematical formulation of the imaging model is provided as follows:

Consider *K* 3D template volumes 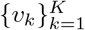, where *v*_*k*_ ∈ ℝ^*b*×*b*×*b*^. Let 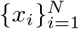 denote the cryo-EM images, where *x*_*i*_ ∈ ℝ^*b*×*b*^. Each image *x*_*i*_ is assumed to be generated from one of the *K* template volumes. Let *c*_*i*_ ∈ {1, …, *K*} represent the structure assignment for *x*_*i*_. Each image *x*_*i*_ is modeled as the result of applying a projective transformation 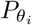 to the template 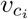, followed by the cryo-EM imaging process *g* and additive noise. Formally,

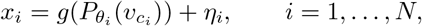

where

- 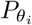 is a projective transformation operator parameterized by *θ*_*i*_ = (*ϕ*_*i*_, *t*_*i*_) ∈ *SO*(3) × ℝ^2^, with *ϕ*_*i*_ and *t*_*i*_ representing the rotation and translation parameters, respectively.
- *g* is the cryo-EM imaging operator (e.g., convolution with the point spread function).
- *η*_*i*_ is a noise term modeling imaging artifacts.

In this work, we aim to obtain the structural assignments *c*_*i*_ ∈ {1, …, *K*} for each 2D cryo-EM image *x*_*i*_.

### 1.2 Our Proposed Method

Our method is built upon an autoencoder framework that learns a mapping from the image space *X* ⊂ ℝ^*b*×*b*^ to a latent space *Z* ⊂ ℝ^*d*^. Specifically, we learn a latent representation **z**_**i**_ ∈ *Z* for each *x*_*i*_ ∈ *X*. We also learn a high-contrast 2D structural template *y*_*i*_ ∈ ℝ^*b*×*b*^ per latent representation *z*_*i*_ using a generator network. The optimal in-plane pose aligning each template *y*_*i*_ with its corresponding input image *x*_*i*_ is estimated through iterative optimization. An in-plane pose is defined as

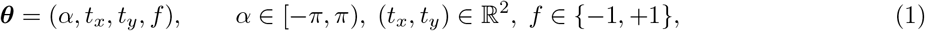

where *α* is the rotation angle, (*t*_*x*_, *t*_*y*_) the translation, and *f* an optional mirror flip.

### 1.3 Input Preprocessing

Before passing the input images to our framework, we perform several preprocessing steps across different stages of our training pipeline. We describe the preprocessing operations as follows:

#### Binning (Shrink)

In Stage 1 of our training, the projection *x* is down-sampled by a factor of 2 using a 2 × 2 average-pooling operator, denoted *S*. Thus, *x*_*s*_ = *Sx*. In later stages of training, the full resolution is kept (*x*_*s*_ = *x*).

#### Low-pass filtering

A fixed isotropic Gaussian kernel of size 3 × 3 is applied via convolution, denoted ℒ. The kernel *g* is defined as:

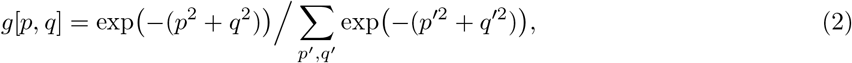

where (*p, q*) are integer coordinates relative to the kernel center. The filtered image is ℒ*x*_*s*_.

#### Circular mask

A smooth circular mask ℳ is applied to suppress boundary artifacts in the target image and the warped template during loss calculation. The mask preserves a central circular region, transitioning smoothly to zero via a cosine ramp. Specifically, for an image of half-size *R*_*h*_ (e.g., *R*_*h*_ = *B/*2 or *B/*4 if binned), the mask has a flat unit value up to radius *R*_*h*_ ™ 4 pixels, followed by a cosine ramp of 4-pixel width.

#### Geometric warp

The differentiable warp corresponding to a pose ***θ***, denoted *T*_***θ***_, transforms an image *x*^′^ as:

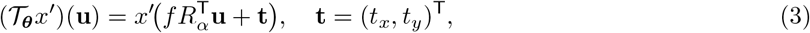

where *R*_*α*_ is the 2D rotation matrix for angle *α*. This operation is implemented using bilinear interpolation.

### 1.4 Network Architecture

Our model architecture consists of an encoder *E* and a decoder *D*, both of which are used across all stages of our training pipeline. For the final stage, we also use a booster decoder *D*^′^ after the decoder. We describe the network architectures of the encoder, decoder, and booster decoder as follows:

#### Encoder *E*

Let *x*_*i*_ be an input projection. After optional shrinking (*S*) and low-pass filtering (ℳ), the result is 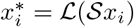 (where S is the identity if not shrinking). The encoder takes as input 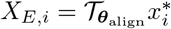, where ***θ***_align_ is an alignment pose (see Sect. 1.5). *X*_*E,i*_ is processed by four 5 × 5 convolutional layers with ReLU activations. These layers are interleaved with 2 × 2 max-pooling operations and dropout layers (rate 0.1). Global max-pooling converts the final feature map to a vector, which then branches into two outputs: *1)* a latent code **z**_*i*_ ∈ ℝR^8^ (further processed with a dense layer and 0.3 dropout); and *2)* a two-element scale–bias vector **m**_*i*_ = (*m*_1_, *m*_2_), derived via dense layers and initialised effectively towards (1, 0).

#### Decoder *D*

Starting from the latent code **z**_*i*_, a fully connected layer expands it to a low-resolution feature map (e.g., *B/*16 × *B/*16 × 8). This map is successively up-sampled by three transposed-convolution blocks (each using 5 × 5 kernels, stride 2, and sine activations, followed by dropout). Two final 5 × 5 convolutional layers (with sine activations, the last one outputting a single channel and scaled by a factor of 10) produce the initial template *D*(**z**_*i*_). The reconstructed template is then affine transformed:

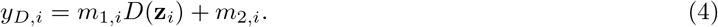

#### Booster decoder *D*^′^

This decoder is active only in Stage 3 of our training pipeline. It takes the feature map *y* _ℒ,*i*_ = ℒ (*y*_*D,i*_) (output of the main decoder after low-pass filtering) as input. The booster consists of a 5 × 5 transposed convolution (stride 2, sine activation, dropout) followed by two 5 × 5 convolutions (sine activations, dropout, with the last layer outputting 1 channel and scaling the output by 10), producing a refined map *y*_refine,*i*_. Concurrently, *y* _ℒ,*i*_ is also upsampled using a 2 × 2 transposed convolution (stride 2, learned kernel) to produce *y*_up,*i*_. The final boosted template *y*_*B,i*_ is a learned linear combination of these two paths and a bias term:

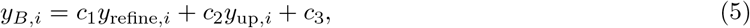

where *c*_1_, *c*_2_, *c*_3_ are learnable scalar coefficients (initialized towards 1, 1, and 0 respectively).

### 1.5 Training Stages

The network is trained in three sequential stages, with weights from stage *s* ™ 1 used to initialize stage *s*. All stages use the loss functions defined in Sect. 1.6, but differ in input resolution, network configuration, and alignment strategy.

**Table 1:** Configuration of the three training stages. ***θ***_align_ refers to the alignment pose used for encoder input and target generation. *T*_fine_ is used for the loss minimization search in all stages.

| Stage | Input Resolution | Encoder Alignment Grid | Booster Head |
|---|---|---|---|
| 1 | Half ($\mathcal{S}x$) | $\mathcal{T}_{\text{coarse-align}}$ | × |
| 2 | Full (*x*) | $\mathcal{T}_{\text{fine-align}}$ | × |
| 3 | Full (*x*) | $\mathcal{T}_{\text{fine-align}}$ | ✓ |

**Stage 1:** The input projections *x*_*i*_ are down-sampled by 2 (*Sx*_*i*_). The encoder input *X*_*E,i*_ and the reconstruction target 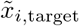 are formed using alignment poses from *T*_coarse-align_ relative to the current estimate. The booster head is inactive. The decoder template *y*_*D,i*_ is low-pass filtered to 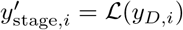.

**Stage 2:** Input projections *x*_*i*_ are at full resolution. The alignment poses for encoder and target are from *T*_fine-align_. The booster head remains inactive. Since the decoder *D* produces a low-resolution map by design, its output *y*_*D,i*_ (after affine scaling and low-pass filtering to ℒ (*y*_*D,i*_)) is upsampled by a learned 2 × 2 transposed convolution *U* and then low-pass filtered again: 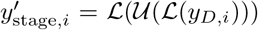. This matches the full resolution of *x*_*i*_.

**Stage 3:** This stage also uses full-resolution inputs *x*_*i*_ and *T*_fine-align_ for initial alignment. The booster decoder *D*^′^ is activated. The template *y*_*D,i*_ is first low-pass filtered, then processed by the booster decoder (Eq. (5)) to yield *y*_*B,i*_. A final low-pass filter is applied: 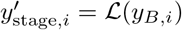.

In all stages, the final per-particle pose is updated iteratively by finding the ***δθ***^*^ from the minimization in Eq. (6) and composing it with the ***θ***_align_ used for that iteration. The candidate alignment poses ***θ***_align_ for the next iteration will be relative to this updated pose.

### 1.6 Training Objective

For each input projection *x*_*i*_, let 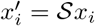 be the (potentially shrunk) version. The network aims to reconstruct a target view of this projection. The target image for the loss calculation is 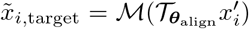,where ***θ***_align_ is the same alignment pose used for the encoder input (see Sect. 1.5). The template generated by the decoder, *y*_*D,i*_ (Eq. (4)), undergoes further processing depending on the training stage to produce

- Stage 1: 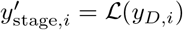
- Stage 2: 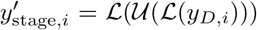,where *U* is a 2 × 2 upsampling (transposed convolution with stride 2, kernel initialized to ones) applied because *y*_*D,i*_ is low-resolution while *x*_*i*_ is full-resolution.
- Stage 3: 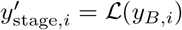, where *y*_*B,i*_ is from Eq. (5).

The reconstruction error is then defined by searching over a finite set *T*_fine_ of candidate refinement poses (Sect. 1.7):

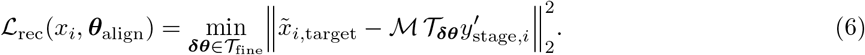

A soft regulariser encourages the latent codes **z**_*i*_ to remain within a sphere of radius *τ* = 20:

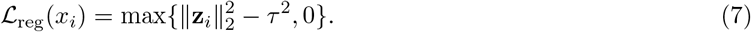

The total per-sample loss is ℒ (*x*_*i*_) = ℒ _rec_(*x*_*i*_, ***θ***_align_) +*λ* ℒ _reg_(*x*_*i*_), with *λ* = 0.01. Network weights are updated using the Adam optimizer. The pose ***θ***_new_ for *x*_*i*_ is updated by composing ***θ***_align_ with the optimal ***δθ***^*^.

### 1.7 Pose Search Grids

Let Δ*α* = 10^°^ and Δ*t* = 2 px define basic angle and translation steps. The discrete pose search grids are:

- *T*_coarse-align_: Used for initial alignment ***θ***_align_ in Stage 1. It consists of rotations *k* · 30^°^ for *k* = 0, …, 11, zero translation (0, 0), and both flips *f* = ±1.
- *T*_fine-align_: Used for initial alignment ***θ***_align_ in Stages 2 and 3. It consists of rotations *k* · Δ*α* for *k* = ™5, …, 5 (i.e., a ±50^°^ range relative to the current estimate), zero translation (0, 0), and no flip (*f* = 1).
- *T*_fine_: Used for the refinement search ***δθ*** in Eq. (6) across all stages. It consists of relative rotations *k* · Δ*α* for *k* = ™5, …, 5; relative translations (*t*_*x*_, *t*_*y*_) where *t*_*x*_, *t*_*y*_ ∈ {0, ±Δ*t*, ±2Δ*t*} (i.e., {0, ±2, ±4} pixels); and no flip (*f* = 1).

The alignment poses ***θ***_align_ are generated by taking the current best estimate for a particle’s pose and applying transformations from *T*_coarse-align_ or *T*_fine-align_. The encoder processes inputs generated from these alignments. The model processes a set of such alignments. One representative alignment from this set is used as the target for the objective (Eq. (6)) during the corresponding training stage.

### 1.8 Particle Grouping and Selection

After training, each particle image *x*_*i*_ is characterized by its latent representation **z**_*i*_ and reconstruction loss ℒ _rec_(*x*_*i*_, ***θ***_align_). Depending on the analysis objective, these outputs are used for particle curation, generation of representative decoded images, and decoded-template-guided particle selection.

#### Particle curation

For the *β*-galactosidase and PKM2 experiments, we concatenated **z**_*i*_ with the corresponding scalar reconstruction loss to form a nine-dimensional feature vector for each particle. The features were standardized and embedded into two dimensions using Uniform Manifold Approximation and Projection (UMAP) [36], after which a two-component Gaussian mixture model (GMM) was fitted to the resulting embedding. The resulting GMM assignments were used to select particle subsets for downstream 3D reconstruction.

#### Generation of representative decoded images

For the TRPV1 experiment, we first retained the 90% of particles with the lowest reconstruction losses. Their latent representations were projected into two dimensions using principal component analysis (PCA) and grouped using K-means with *K* = 25. The cluster centers were mapped back to the latent space and decoded to generate representative decoded images. For the TRPV1 and *β*-galactosidase mixture, representative locations in the latent-space representation were also decoded to examine the separation of the two particle classes.

#### Decoded-template-guided particle selection

For the *Chlamydomonas* cryo-ET experiment, we used the reconstruction loss to first filter particle images and then embedded the latent representations **z**_*i*_ of the retained particles into two dimensions using UMAP. We sampled the resulting embedding on a 20 × 20 regular grid and decoded the corresponding latent representations to generate representative 2D structural templates. Regions containing distinct structural patterns were manually selected, and the corresponding particle subsets were mapped back to the original subtomograms for downstream 3D analysis.

### 1.9 Conventional Classification and Reconstruction

We used conventional cryo-EM/ET classification and reconstruction methods in two experimental settings. For the *Chlamydomonas* experiment, cryoSPARC [30] 2D classification served as a conventional baseline for particle grouping, and particle subsets derived from the classification results were subsequently used for cryoSPARC ab-initio reconstruction. We additionally applied RELION 3D classification [31] directly to the corresponding 3D subtomograms. For the single-particle cryo-EM experiments, cryoSPARC ab-initio reconstruction was used to compare the reconstruction quality obtained using all particles with that obtained after particle curation by our method.

#### 2D classification

cryoSPARC 2D classification groups particle images into a specified number of classes while estimating their in-plane rotations and translations. Using expectation–maximization, particle poses and probabilistic class assignments are iteratively updated, and the aligned particle images are weighted to update the corresponding class averages. These class averages provide visual summaries of the classes and can be used to select the corresponding particle subsets for subsequent 3D reconstruction.

#### Ab-initio reconstruction

cryoSPARC ab-initio reconstruction uses likelihood-based optimization to estimate a specified number of 3D structures from the observed particle images. Starting from randomly initialized volumes, it iteratively estimates particle poses and structural assignments and updates the volumes using stochastic gradient descent.

#### 3D classification

RELION 3D classification separates particles into a specified number of structural classes using a regularized likelihood framework. During optimization, the class probabilities and corresponding 3D class volumes are iteratively updated.

## Data availability

All data used in the paper are publicly available through EMPIAR. Structures produced in this paper will be deposited in the EMDB upon paper acceptance.

## Code availability

The source code for the method described in this study is available at https://github.com/Qifeng-Wu99/cryo2dclass.

## Acknowledgments

This work was supported in part by U.S. NIH grant R35GM158094. This research is also supported by NIH grant R01GM150905 (M.C.) and U.S. DOE under BCIS FWP-101359 (M.C.).

